# The degree of T cell stemness differentially impacts the potency of adoptive cancer immunotherapy in a Lef-1 and Tcf-1 dependent manner

**DOI:** 10.1101/2023.03.08.531589

**Authors:** Guillermo O. Rangel Rivera, Connor J. Dwyer, Hannah M. Knochelmann, Aubrey S. Smith, Arman Aksoy, Anna C. Cole, Megan M. Wyatt, Jessica E. Thaxton, Gregory B. Lesinski, Chrystal M. Paulos

## Abstract

Generating stem memory T cells (T_SCM_) is a key goal for improving cancer immunotherapy. Yet, the optimal way to modulate signaling pathways that enrich T_SCM_ properties remains elusive. Here, we discovered that the degree to which the PI3Kδ pathway is blocked pharmaceutically can generate T cells with differential levels of stemness properties. This observation was based on the progressive enrichment of transcriptional factors of stemness (Tcf-1 and Lef-1). Additional investigation revealed that T cells with high stemness features had enhanced metabolic plasticity, marked by heightened mitochondrial function and glucose uptake. Conversely, T cells with low or medium features of stemness expressed more inhibitory checkpoint receptors (Tim-3, CD39) and were vulnerable to antigen-induced cell death. Only TCR-antigen specific T cells with high stemness persisted following adoptive transfer *in vivo* and mounted protective immunity to melanoma tumors. Likewise, the strongest level of PI3Kδ blockade *in vitro* generated human tumor infiltrating lymphocytes (TILs) and CAR T cells with heightened stemness properties, in turn bolstering their capacity to regress human mesothelioma tumors. We find that the level of stemness T cells possess *in vitro* differentially impacts their potency upon transfer in three tumor models. Mechanistically, both Lef-1 and Tcf-1 sustain anti-tumor protection by high T_SCM_, as deletion of either one compromised cellular therapy. Collectively, these findings highlight the therapeutic potential of carefully modulating PI3Kδ signaling in T cells to confer high stemness and mediate protective responses to solid tumors.

## Introduction

The quality of the T cells generated for adoptive cell transfer (ACT) therapy for solid tumors impacts clinical outcome (*1*–*4*). Several features related to the “youth” of T cells are associated with greater clinical efficacy in patients (*5*, *6*). T cells with stemness properties have emerged as ideal subsets for immunotherapy–termed stem-like memory T cells (T_SCM_) (*7*–*9*). T_SCM_ self-renew, give rise to effector cells with polyfunctional properties and mediate durable anti-tumor responses (*10*). Patients responsive to immune checkpoint blockade therapy also have more T_SCM_ in the tumor (*11*, *12*). Many approaches have been employed to modulate T cell differentiation, including enriching T_SCM_ with cytokines or pharmacologic agents (*13*–*16*). However, how to properly generate and identify T cells with durable features of stemness and potent antitumor activity remains underexplored.

We hypothesized that T cells can be generated to exhibit differential properties of stemness. We also anticipate that T cells with the greatest stemness properties would be more effective at clearing tumors. To test this idea, we inhibited the p110δ subunit of the PI3K signaling pathway (i.e., PI3Kδ), as this subunit regulates their differentiation. Using escalating doses of a PI3Kδ inhibitor idelalisib, we generated T cells with three degrees of stemness, marked by their low, medium, to high Lef-1 and Tcf-1 expression. T cells expanded in the absence of this drug possessed an exhausted progenitor-like phenotype (T_EP_). T cells with the highest stemness properties were bioenergetically plastic, able to use glucose and sustain mitochondrial mass and function even after tumor rechallenge. Moreover, only T cells with high degrees of stemness resisted acquisition of markers related to T cell exhaustion and mediated long-lived immune responses in mice with melanoma. Likewise, human TIL and mesothelin specific CAR T cells could be bestowed with progressive levels of stemness via PI3Kδ blockade. CAR T cells with high stemness mediated robust immune responses against human mesothelioma. Mechanistically we found that high T_SCM_ mediate antitumor responses dependent on Lef-1, regardless of Tcf-1 expression. Our data suggests a nonredundant role for Lef-1 in antitumor T_SCM_ activity. This work highlights for the first time that T cells with differential stemness levels can be generated by targeting PI3Kδ signaling and identify distinct metabolic and transcription properties that can be used to select potent T_SCM_ cells to treat cancer.

## Methods and Materials

### Animals

Pmel-1 transgenic mice, male and female, were obtained from The Jackson Laboratory, and maintained as homozygous colony. For all studies Pmel-1 were sacrificed between 7 to 10 weeks after birth. C57BL/6J female mice for *in vivo* studies were obtained from The Jackson Laboratory and inoculated with tumors subcutaneously at 7 to 10 weeks of age. C57BL/6J mouse studies were housed at Emory’s pathogen free (SPF) DAR animal facility in clinic B, and Whitehead Institute. NSG from Jackson Laboratories mouse studies were performed at the Medical University of South Carolina at their pathogen free clean facility.

### T cell cultures

Spleens from Pmel-1 transgenic mice were processed through 70μm filters, RBC lysed for 5 min at room temperature and then resuspended in complete media, RPMI 1640 medium supplemented with 2mM L-glutamine, 1X non-essential amino acids, 1mM HEPES, 50U/mL penicillin, 50μg/mL streptomycin, and 10% heat-inactivated fetal bovine serum (Atlas biologicals). Splenocytes were activated with human glycoprotein 100 (hgp100) peptide at a final concentration of 1μM in the presence of recombinant human 100IU/mL IL-2 (NIH repository), then plated at a density of 1×10^6^/mL in 2mL of complete media in a 24-well plate and placed in a 5% CO_2_ incubator at 37 C. Titration of idelalisib (0.1 μM, 1 μM or 10μM) or DMSO control was added to media during T cell expansion and then phenotype assessed after 7 days in culture.

### Adoptive T cell therapy protocol

Melanoma B16F10 (containing either a mouse low affinity EGS or human high affinity KVP epitope of glycoprotein 100) were washed in sterile PBS twice and injected subcutaneously into the abdomen of B6 mice at 5×10^5^cells/mouse in 100μL. One day prior to T cell infusion, mice were lymphodepleted using 4Gy total body irradiation (TBI). Pmel-1 T cells expanded with escalating doses of idelalisib as noted for one week, they were washed off from the drug re-stimulated and then infused into mice. IL-2 complex was made with 1.5μg of recombinant human IL-2 (NIH) and 7.5μg of anti-human IL-2 (R&D Systems) per injection in 100μL of sterile PBS. Three doses of IL-2 complex were administered post ACT every other day starting on the day of T cell infusion. Tumor area was measured twice a week until endpoint criteria was met. Tumor analysis of T cell infiltration and phenotype was done by mincing melanoma tumors and incubating in DNase I and Collagenase IV in Hank’s Buffered Saline (HBS) for 1hr at 37 °C. Tumor samples were then filtered in 70μm screens and then passed a second time over 70μm mesh 17×100mm 14mL tubes.

### Mesothelin CAR T cell production and ACT

Bulk CAR T cells were activated with CD3/CD28 bead (1 T cell: 1 Bead) ratio for 3 days, then debeaded and expanded for 10 days with or without 0.1μM, 1μM or 10μM idelalisib and given 200 IU/mL of IL-2 throughout *in vitro* expansion. T cell transduction efficiency was on average 35-42.5% for Mesothelin specific CAR, transferred CAR T cell number was not adjusted to mesothelin positive percentage. M108 cell lines were expanded nearly 3 weeks *ex vivo* and then NSG mice were inoculated subcutaneously and allowed to grow for 48 days. 2×10^6^ CAR T cells were infused on day 48 and tumor measurements were taken twice a week until tumor endpoints were met.

### Lung metastatic burden analysis

Lungs and draining lymph nodes were then processed like spleen as described above. For tumor lung burden ImageJ was used to measure the area covered by melanoma compared to normal lung and then plotted as percentage of lung area occupied by melanoma per lung lobe.

### Flow cytometry

Single-cell suspensions were generated and prepared in FACS buffer, and then cells were stained with antibodies for surface markers for 15 min at room temperature in the dark. Intracellular cytokine stain was performed following restimulation of pmel-1 T cells as above with the addition of 1X Brefeldin A (Biolegend) and 1X Monensin (Biolegend) following incubation period of 4hrs in both unstimulated and restimulated groups prior to analysis. For transcription factor staining T cells and processed tissue were fixed with eBioscience™ Foxp3 / Transcription Factor Staining Buffer Set (ThermoFischer) according to the manufacturer. Phenotypic characterization was performed using the FACS Verse or Symphony A3/A5 (BD Biosciences).

### Plate bound re-stimulation

Pmel-1 CD8^+^ T cells were activated with hgp100 and cultured in the presence or absence of idelalisib for one week. After one week of expansion, T cells were collected, washed from compound containing media and plated into 30ng/mL CD3ε coated 48-well plates and incubated over 24hr. Cells were stained for Annexin V and PI at room temperature for 30min in Annexin V buffer at a dilution of 1:1000 and then assessed by flow cytometry.

### CRISPR/Cas9 gene editing

Lyophilized crRNAs targeting *Tcf7* and *Lef1* and tracrRNA were chemically synthesized (IDT) and resuspended in nuclease-free duplex buffer at a stock concentration of 200μM. A *Rosa26* CRISPR crRNA guide was synthesized separately by Synthego as a CRIPSR-Cas9 knockout control as previously described (*17*). RNP complexes for three different guide RNAs were then pooled into one reaction. Immediately after electroporation, cells were incubated at 37 °C in media saved from initial 24hr activation and drug treatment. Knockout efficiency was measured 48 hours and 7 days post-electroporation by flow cytometry and qPCR.

### Metabolism assays

Oxygen consumption rate (OCR) measurements either 2.5×10^5^ or 1×10^6^ pmel-1 cultured T cells were used for the 96-well or 24-well plate format for the seahorse bioanalyzer, respectively. T cells were seeded onto plates using Cell-Tak and mitochondrial stress test was measured in XF media as per manufacturer. For glucose uptake assay, T cells were cultured for 30min at 37C° in glucose free complete media in the presence of 1:200 2-NBDG, following incubation T cells were stained with surface antibodies and viability dye zombie aqua or zombie UV. Mitochondrial dyes to measure mass and potential were used as following tetramethyl rhodamine methyl ester (TMRM) was incubated at 250nM for 30mins at 37 C° in complete media prior to flow cytometry staining, and Mitotracker FM deep red was stained at 20nM along with surface flow cytometry stains at room temperature (both Thermo Fischer Scientific).

### Western blotting

10×10^6^ T cells were resuspended in 100μL RIPA buffer containing Halt© protease and phosphatase inhibitor (ThermoFisher). 10μg of protein was loaded into 4-20% Tris Glycine pre-cast gels. Protein was then transferred into a PVDF membrane. The PVDF membrane was then blocked in 5% skim milk Tris buffered saline 0.1% Tween20 buffer. Membrane was probed for primary antibodies as listed in supplemental table. SuperSignal™ West Pico PLUS Chemiluminescent Substrate (ThermoFisher) was used to develop signal. Bio-Rad ChemiDoc MP Imaging System was used to detect chemiluminescence. Relative optical density and analysis of bands of interest was performed using the Image J.

### RNA sequencing

Pmel-1 T cells activated with 1μM hgp100 and cultured with 0, 0.1μM, 1μM and 10μM idelalisib for 7 days. T cell media was washed thoroughly to get rid of inhibitor and then restimulated with 10Gy irradiated B6 splenocytes loaded with 1μM hgp100 at a ratio of 1 splenocytes to 10 T cells, then cultured for 24hr hours. On Day 8 all 5 biological replicates and 4 unstimulated and 4 restimulated conditions were then taken (total of 40 samples) at 10×10^6^ T cells for processing with Trizol to isolate RNA for bulk RNA-sequencing.

### Human TIL

Resected lymph node biopsies with metastatic lesions were collected after deidentification and incubated with complete media containing antimycin overnight prior to processing. Tissues were then cut into ~3mm diameter pieces and plated into individual wells in a 24-well plate containing complete media with 6,000 IU IL-2/mL and given 30ng/mL OKT-3 CD3ε agonist antibody. Cultures were maintained for 5 days until T cells egressed, then idelalisib was added as in the mouse cultures as follows: low dose (0.1μM), medium dose (1μM), and high dose (10μM). Human TIL was then analyzed 14-21 days after expansion.

### Statistical analysis

Boxplots show mean of data and interquartile with outliers, bars show either ±SD or ±SEM as noted in caption. Non-paired t test (two tailed) was used for the comparison of 2 datasets. ANOVA was used for > 2 comparison groups, and comparisons across groups was performed with Bonferroni post-hoc correction. For survival comparisons Mantel-Cox regression were performed between groups.

### Ethics approval

All human tissue samples were received in a deidentified manner from the Emory from Cancer Tissue and Pathology shared resource of Winship Cancer Institute of Emory University and NIH/NCI under award number P30CA138292. All animal procedures performed at the Medical University of South Carolina or Emory University were approved by each university’s Institutional Animal Care & Use Committee, protocol number 0488 or 201900225, respectively.

### Data and materials availability

All data associated with this study are present in this paper or the Supplemental materials.

## Results

### Escalating PI3Kδ inhibition progressively enriches T cells with accentuated stemness properties

We reported that the anti-tumor activity of adoptively transferred T cells is improved when *in vitro* conditioned with idelalisib, which inhibits the phosphatidylinositol-3-kinase δ (PI3Kδ) signaling pathway (*18*–*20*). We posited that full blockade of the PI3Kδ pathway is needed to manufacture cell products with optimal stem-like memory properties. To test this idea, pmel-1 CD8^+^ T cells (*which express a transgenic TCR that recognizes the antigen glycoprotein 100 (gp100) expressed on melanoma*) were expanded for one week in the presence of escalating idelalisib concentrations, ranging from 0.1μM (low), 1μM (medium) to 10μM (high) in the presence of 100IU/mL of IL-2 (**Fig. 1A**). As expected, increasing idelalisib concentration progressively reduced T cell differentiation, as marked by a distinct decrease in CD44 and increase in CD62L expression (**Fig. 1B**). Conversely, 40% of uninhibited T cells displayed an effector-like profile (CD44^+^ CD62L^-^). However, high PI3Kδ blockade could enrich a large proportion of CD44^-^ CD62L^+^ T cells *in vitro*. We consistently observed the generation of these distinct T cell populations in more than 15 independent experiments (**Fig. 1C**).

**Figure 1.**
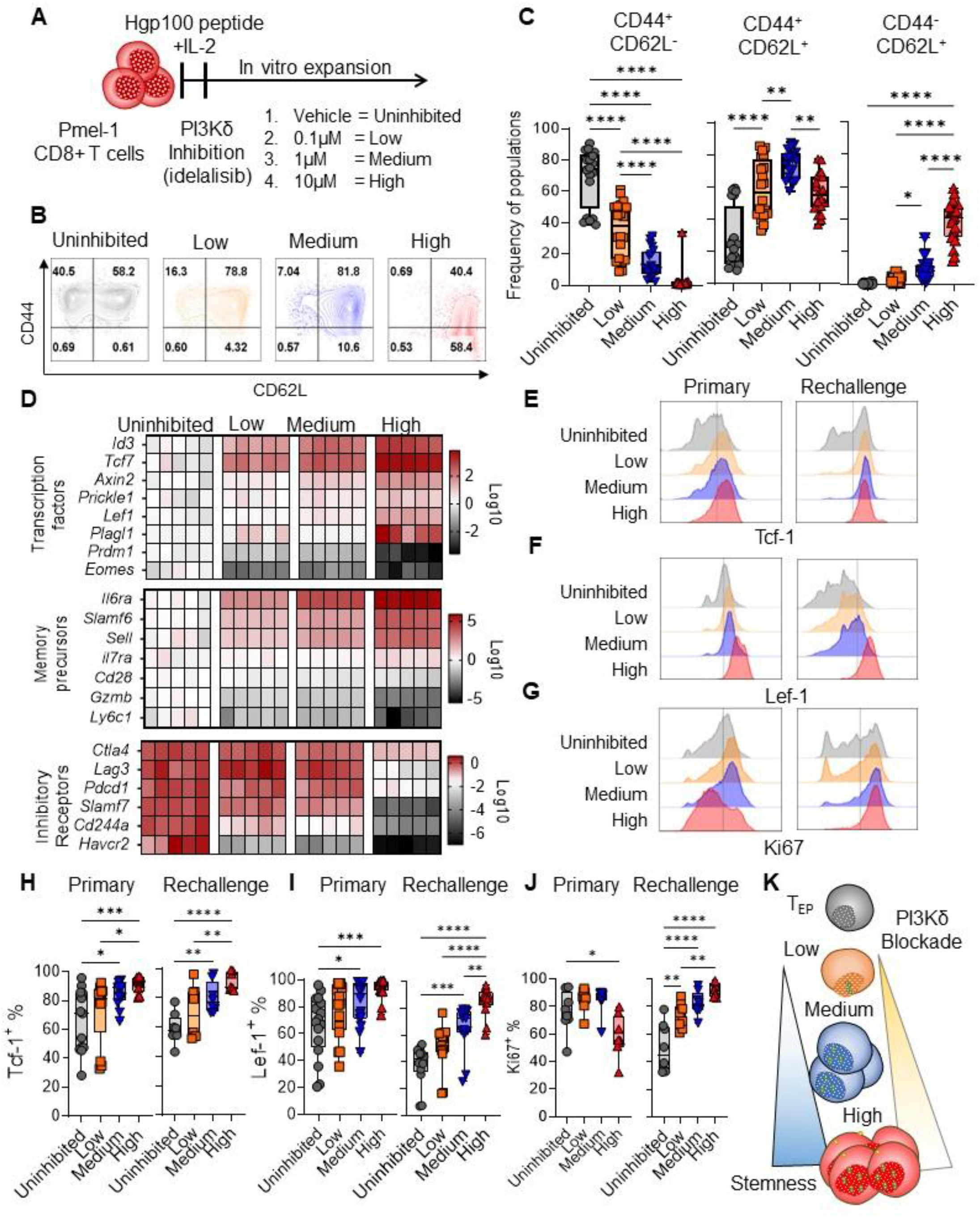
Progressive PI3Kδ inhibition step-wise enriches CD8^+^ T cells with increased features of stemness. **A)** Pmel-1 T cell splenocytes were TCR-activated with 1μM hgp100 and given 100IU/mL IL-2 and then treated with indicated amount of PI3Kδ-inhibitor idelalisib 3 hours later. Phenotype of pmel-1 T cells was then assessed after 7 days of expansion *in vitro*. **B**) T cell differentiation based on CD44 and CD62L flow cytometry contour plots, populations defined as CD44 single positive (SP), CD44 CD62L double positive (DP), and CD62L single positive (SP). **C**) Quantitation of B. N=20 distinct cultures per condition, representative of 5 different experiments. **D**) Bulk RNA-seq for progressively PI3Kδ-inhibited pmel-1 T cells expanded for 7 days, showing select genes related to stemness signatures, data is log2 transformed, the fold change is expression normalized from uninhibited pmel-1 T cells. N=5 per condition. *Primary versus tumor rechallenge:* Pmel-1 T cells expanded for 7 days were washed to get rid of the inhibitor, then reactivated with tumor antigen by co-culturing them with 10Gy irradiated B6 splenocytes pulsed with 1μM hgp100 for 24hr, a procedure that recapitulate tumor antigen rechallenge *in vitro*. Expression of **E**) Tcf-1, **F**) Lef-1 and **G**) Ki67 during primary expansion and secondary antigen rechallenge. **H**) Quantitation of data in E, **I**) F, and **J**) G, respectively. N=6-13 mice per group, representative of 3-6 independent experiments. **K**) Visual classification of stemness grades acquired by T cells with serial PI3Kδ inhibition with idelalisib based on transcriptional signatures of T cell stemness, expression of Tcf-1 and lef-1, and proliferative potential before and after tumor antigen rechallenge. ANOVA analysis with Bonferroni post-hoc correction for multiple comparisons. ρ≤0.05, * p≤0.05, ** p≤0.01, *** p≤0.001, **** p≤0.0001 were considered significant, while ρ values >0.05 were considered not significant (ns).

### PI3Kδ blockade enriches unique transcriptional signatures of T cell stemness

Genome-wide RNA-seq analysis revealed that progressive PI3Kδ blockade imparted T cells with unique gene signatures associated with T cell stemness. As shown in F**ig. 1D**, T cells expanded with increasing idelalisib concentrations enriched transcription regulators (*Tcf7, Lef1, Plagl1, Axin2, and Id3*) and surface markers (*Il7ra, Slamf6, Sell and Cd28*) involved in T cell stemness. Concomitantly, transcriptional regulators of T cell effector differentiation (*Prdm1, Eomes*) and inhibitory checkpoint receptor (*Lag3, Pdcd1, Havcr2, Cd244*) were lower in T cells progressively blocked of the PI3Kδ signaling pathway. Other stemness/differentiation markers, including Slamf6, CD28 and Granzyme B, were also distinct among the 4 subsets at the protein level, which matched our RNA seq data set (**Fig. S1 A-B**).

### The level of T cell stemness correlate with Lef-1/Tcf-1 expression, proliferation and cytokine production

We next assessed if idelalisib expanded T cells functioned like stem-like memory cells, which express Tcf-1 and Lef-1, proliferate, and co-secrete cytokines (*21*, *22*). Tcf-1 and Lef-1 were elevated in all T cells treated with any idelalisib amount compared to uninhibited T cells. Yet, following tumor antigen challenge (in absence of idelalisib), both Tcf-1 and Lef-1 were diminished in T cells inhibited with low and medium idelalisib concentration. However, the highest dose of this drug best sustained Tcf-1 and Lef-1 (**Fig. 1E-F, 1H-I**). We also found that high concentration of idelalisib stunted T cell proliferation during the primary expansion. Yet, upon antigen rechallenge, these cells proliferated robustly while uninhibited T cells proliferated poorly (**Fig. 1G, J**). We also found that uninhibited, low and medium PI3Kδ inhibited T cells co-secreted more inflammatory cytokines IFN-γ and TNF-α than high dose PI3Kδ inhibition T cells post-antigen encounter (**Fig. S2A-B**). Further, granzyme B, a cytokine classically produced by effector progenitors (*23*), was mainly produced by low or medium PI3Kδ-inhibiited T cells (**Fig. S2C-D**). Based on their molecular and functional profiles we henceforth term PI3Kδ-treated T cells with their stemness grade as follows: Uninhibited T cells as progenitors of effectors/exhausted T cells (T_EP_), and T cells progressively blocked of PI3Kδ signaling as low, medium and high T_SCM_ cells (shown visually in **Fig. 1K**).

### T cell grades of stemness promote differential mitochondrial function

Metabolic adaptations correlate with T cell differentiation (*24*–*26*). These adaptations shift T cell metabolism from a reliance on anaerobic glycolysis and glutaminolysis (*27*, *28*) to a dependance on mitochondrial metabolism and lipid-based catabolism (*29*, *30*). Further, memory T cells have enhanced mitochondrial respiration (*31*). Thus, we hypothesized that oxygen consumption rate (OCR), specifically spare respiratory capacity (SRC), would be elevated in stem-like memory T cells (T_SCM_) with high degrees of stemness. Although maximal respiration increased commensurably with stemness features (**Fig. 2A**), SRC was most induced in medium and high TSCM (**Fig. 2B)**. Because of the canonical association of FAO and T cell memory development, we surveyed genes involved in this pathway (*32*). To our surprise, FAO transcripts were suppressed in high T_SCM_ and but not in low and medium T_SCM_ (**Fig. 2C**). We also found that mitochondrial potential (TMRM) was robustly induced in high T_SCM_ compared to TEP, low and medium TSCM after primary expansion, and only sustained at high levels in high T_SCM_ (**Fig. 2D, G**). Further, after tumor antigen rechallenge, mitochondrial content was nearly 3-fold greater in high T_SCM_ compared to other subsets (**Fig. 2E, H**). As high mitochondrial membrane potential is related to increased mitochondrial reactive oxygen species (ROS_mt_), we wondered if T cells with high stemness produced more ROS_mt_. Following antigen rechallenge we found that TEP take up high amounts of MitoSox Red, a surrogate for superoxide radicals in the mitochondria. Low and medium T_SCM_ also produced ROSmt, but to a lesser extent than T_EP_ cells. Remarkably, high T_SCM_ produced nominal amounts of ROSmt (**Fig. 2F, I)**. This data collectively suggested that differential stemness states can be defined based on mitochondrial fitness, which we define as T cells that sustain high mitochondrial potential, low mitochondrial ROS, and high mitochondrial content, even after antigen rechallenge.

**Figure 2.**
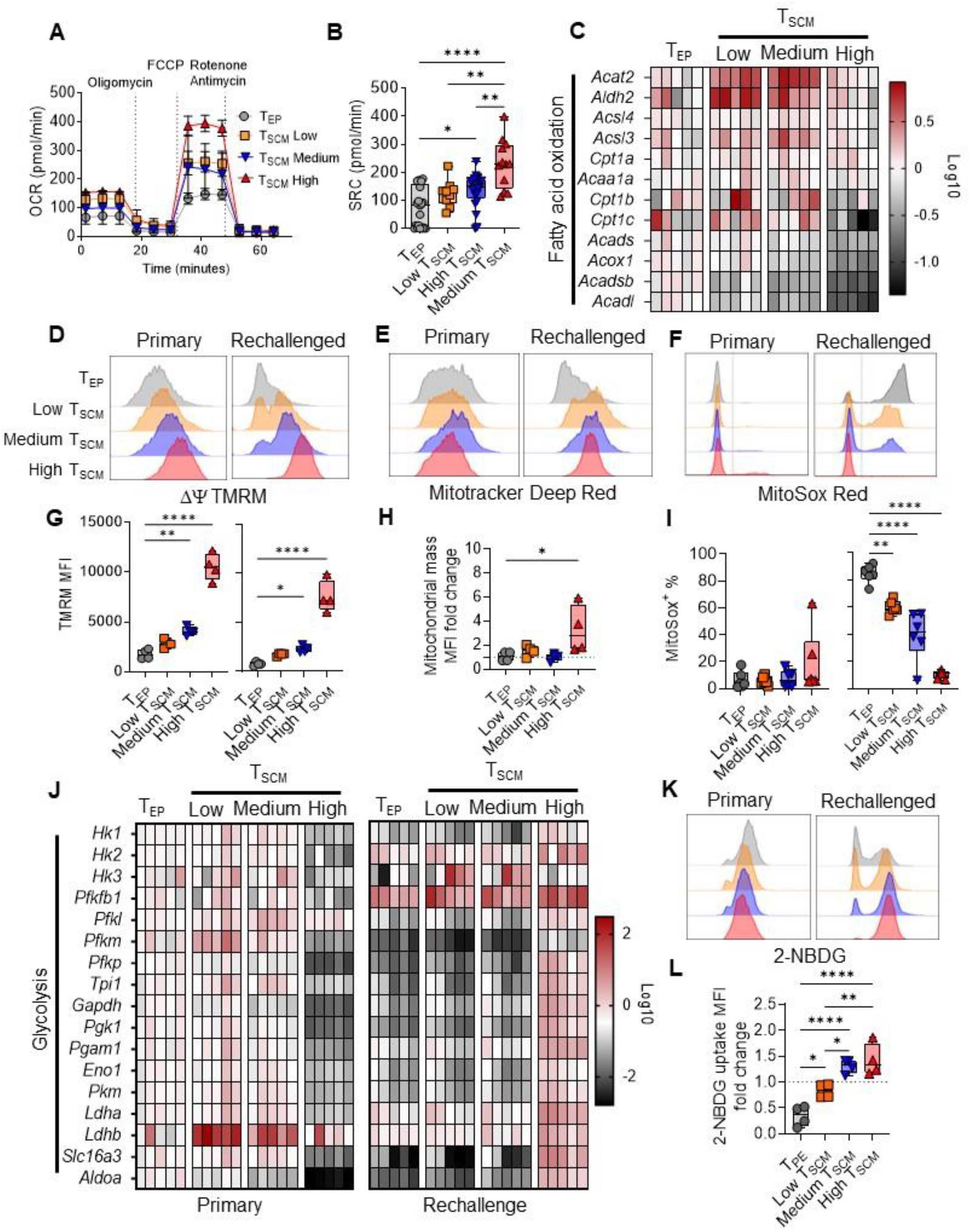
CD8^+^ T cell with the highest stemness have the most bioenergetic plasticity upon tumor recall. As in Fig 1A, pmel-1 stemness subsets were expanded for one week and then reactivated with tumor antigen, as detailed in Fig 1E-L **A**) Seahorse mitochondrial stress test of idelalisib generated T_SCM_ subsets compared to T_EP_. **B**) Spare respiratory capacity, maximal-basal respiration. N=3, 4-5 technical replicates per T_SCM_ subsets, representative of 3 experiments. **C**) RNA-seq genes in the fatty acid oxidation pathway, data is log2 transformed, the fold change is expression normalized from uninhibited pmel-1 T cells. N=5 per condition. **D**) Flow cytometry histograms of mitochondrial dyes for membrane potential (TMRM) and **E**) mitochondrial mass (Mitotracker Deep Red) assessed before and after tumor antigen restimulation for 24 hours. **F**) MitoSox Red (Singlet Oxygen radical surrogate). **G**) Quantitation of median fluorescence intensity (MFI) from D, **H**) fold change MFI from primary and rechallenged T cells in E. **I**) Reactive oxygen species measured by percent MitoSox Red positive populations in F before and after antigen rechallenge. N=6 per condition, representative of 3 independent experiments. **J**) RNA-seq gene list of glycolytic genes during primary expansion and after antigen rechallenge. Data is log2 transformed, the fold change is expression normalized from primary expanded T_EP_ T cells. N=5 per condition. **K**) Flow cytometry histograms of T cells at the end of culture (primary) or after antigen rechallenge (rechallenged), and then assessed for their ability to take up 2-NBDG on day 8. **L**) Median fluorescence intensity (MFI) fold change of 2-NBDG uptake from primary expansion on T cell groups following antigen rechallenge. N=6 per condition, representative of 3 independent experiments. ANOVA analysis with Bonferroni post-hoc correction for multiple comparisons. ρ≤0.05, * p≤0.05, ** p≤0.01, *** p≤0.001, **** p≤0.0001 were considered significant, while ρ values >0.05 were considered not significant (ns).

### High T_SCM_ have heightened glucose usage capacity after tumor antigen exposure

To explore how T cells with distinct stemness engage in glycolysis, we next evaluated these pathways via molecular and functional assays. High T_SCM_ expressed lower levels of glycolytic genes than other subsets, including T_EP_, low or medium T_SCM_ cells (**Fig. 2J left panel**). However, suppression of glycolytic genes in high TSCM was transient, as they were vastly induced following antigen rechallenge compared to other subsets (**Fig. 2J right panel)**. In fact, after antigen rechallenge, high T_SCM_ robustly took up 2-NBDG, a surrogate for glucose uptake (**Fig. 2K)**. Compared to the primary expansion, 2-NBDG uptake was 2-fold greater in both medium and high T_SCM_ compared to low T_SCM_. Conversely, T_EP_ were impaired in glucose uptake (**Fig. 2L),** indicating a diminished plasticity in glucose utilization upon tumor antigen reencounter.

### High T_SCM_ are resistant to tumor antigen-induced-cell death and upregulate Bcl-2

T_SCM_ cells self-renew, give rise to effectors and upregulate anti-apoptotic molecules (*21*, *33*). Thus, we posited that high T_SCM_ may resist cell death following TCR stimulation. Indeed, as expected, post CD3 stimulation, high T_SCM_ remained healthy (59.2% Annexin-PI), as in **Fig. 3A-B**. Conversely, low and medium T_SCM_ rapidly transitioned towards early (Annexin V^+^PI^-^) and late apoptotic states (Annexin+ PI+). Additionally, high T_SCM_ expressed more Bcl-2 prior to antigen exposure (**Fig. 3C-D)**. Collectively, our data suggest that even though T cells can be generated with different stemness levels, their resistance to AICD post antigen recall responses *in vitro* is different. This data may imply that T cells with different stemness levels might mediate varied responses to tumors post ACT.

**Figure 3.**
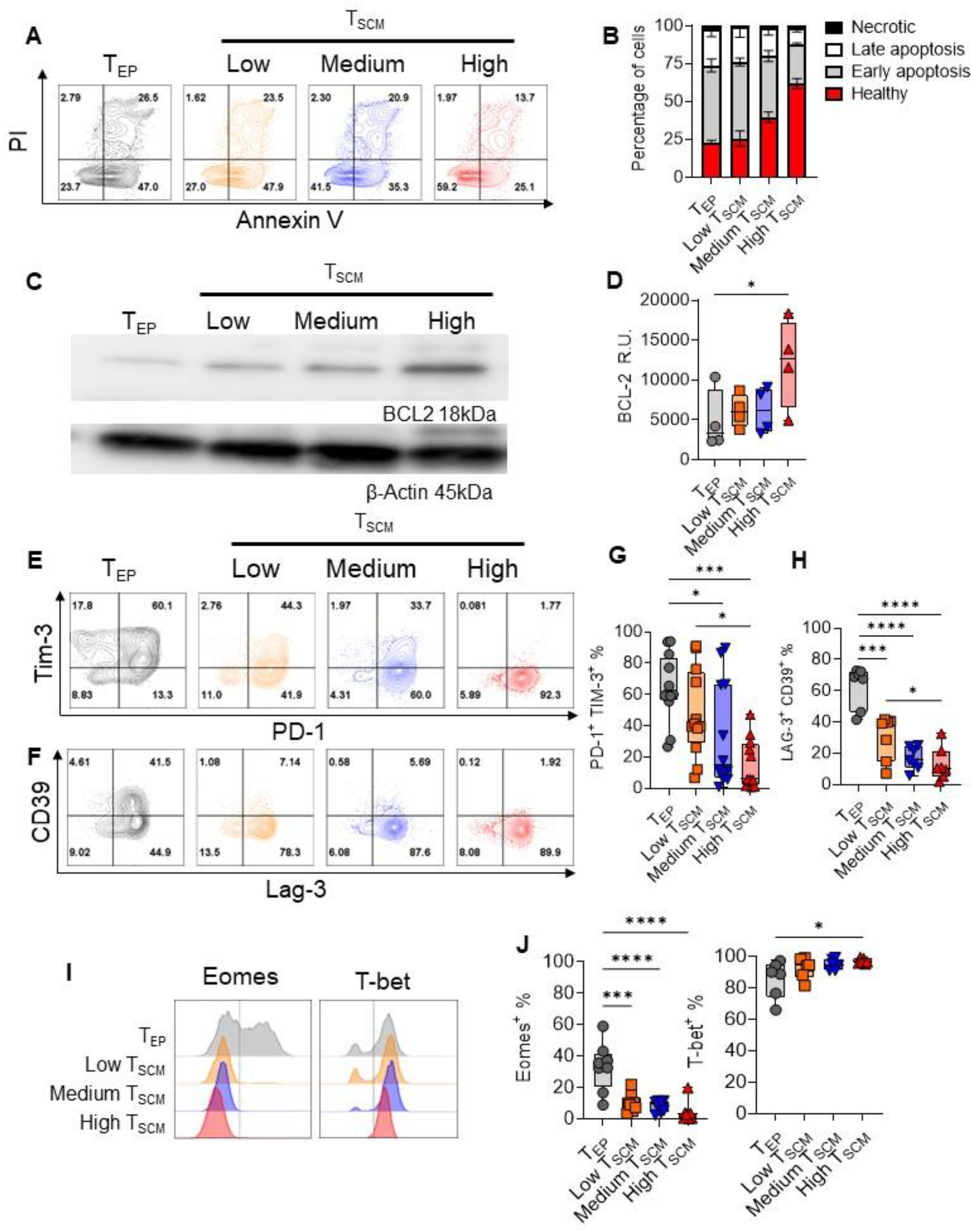
T_SCM_ with high stemness resist cell death and oppose transitioning into an exhausted profile. **A**) Representative flow plots of day 7-T_SCM_ generated with PI3Kδ inhibition pmel-1 CD8+ T restimulated with CD3 coated plates and then assessed for staining of Propidium Iodide (PI) and Annexin V 24 hours later. **B**) Stacked bar graph showing percentage of transitioning populations based on PI and Annexin V. Healthy (PI-Annexin V-), early apoptosis (PI-Annexin V+), late apoptosis (PI+Annexin V+) and necrotic cells (PI+Annexin V-). N=6 per condition, representative of two independent experiments. **C**) Representative blot expression of β-Actin and BCL-2 in induced T_EP_ and various T_SCM_ subsets 7 days after primary expansion. **D**) Relative quantitation normalized to β-Actin. N=6 per group, representative of 3 independent experiments. **E**) pmel-1 CD8^+^ T cells were washed from media containing idelalisib and then re-stimulated for 24hr and then assessed for the expression of Tim-3, PD-1, **F**) Lag-3 and CD39. **G** and **H**) Quantitation on the double positive populations for both E) PD-1+ Tim-3+ and F) Lag-3+ CD39+, respectively. N=7-12 per condition, representative of 4-5 independent experiments. **I**) Histogram expression of Eomes and T-bet on re-stimulated T cell subsets after 24hr hours. **J**) Quantitation of positive populations from I. N=8 per condition, representative of 4 independent experiments. ANOVA analysis with Bonferroni post-hoc correction for multiple comparisons. * p≤0.05, ** p≤0.01, *** p≤0.001, **** p≤0.0001 were considered significant, while ρ values >0.05 were considered not significant (ns).

### Inhibitory checkpoint receptors and transcriptional regulators of exhaustion are suppressed in TSCM with high degrees of stemness

Some T cells in the tumor become exhausted progenitors, defined by transcriptional regulators (T-bet, Eomes, and Tcf-1) and inhibitory checkpoint receptors (e.g., PD-1, Lag-3, Tim-3 and CD39) (*34*–*36*). As expected, Tim-3 was most elevated on T_EP_ compared to low, medium and high T_SCM_ following antigen rechallenge (**Fig. 3E**). PD-1 and Lag-3 however were highly expressed on all subsets. Additional investigation revealed that T_EP_ expressed far more CD39 compared to T_SCM_ subsets (**Fig. 3F**). Notably, few high T_SCM_ co-expressed PD-1^+^ Tim-3^+^ and Lag-3^+^ CD39^+^ compared to other subsets (**Fig. 3G, H**) Finally, we observed that Eomes was lower on all T_SCM_ compared to T_EP_, but lowest in high T_SCM_ while T-bet was slightly enhanced in high T_SCM_ **(Fig. 3I-J).**

### High T_SCM_ mediate potent anti-tumor responses *in vivo*

We hypothesized that T cells with the greatest degree of stemness would mediate the most durable anti-tumor efficacy when infused into mice. To test this idea, we transferred T_EP_, low, medium or high T_SCM_ Thy1.1 pmel-1 CD8^+^ T cells into Thy1.2 mice bearing poorly immunogenic B16F10 melanoma tumors (**Fig. 4A**). Indeed, mice survived long-term when infused with high T_SCM_, resulting in cures in 30% of the animals (**Fig. 4B-C)**. High T_SCM_ engrafted in the peripheral blood most robustly into tumor-bearing mice after 14 days following infusion (**Fig. 4D**). Interestingly, Tcf-1 was partially sustained in high T_SCM_ cells in the primary melanoma and in the tumor draining lymph node (**Fig. 4E-F).** While high T_SCM_ can mediate potent primary responses in animals, it remained unknown if they retained the capacity to mediate long-lived protective responses to tumors.

**Figure 4.**
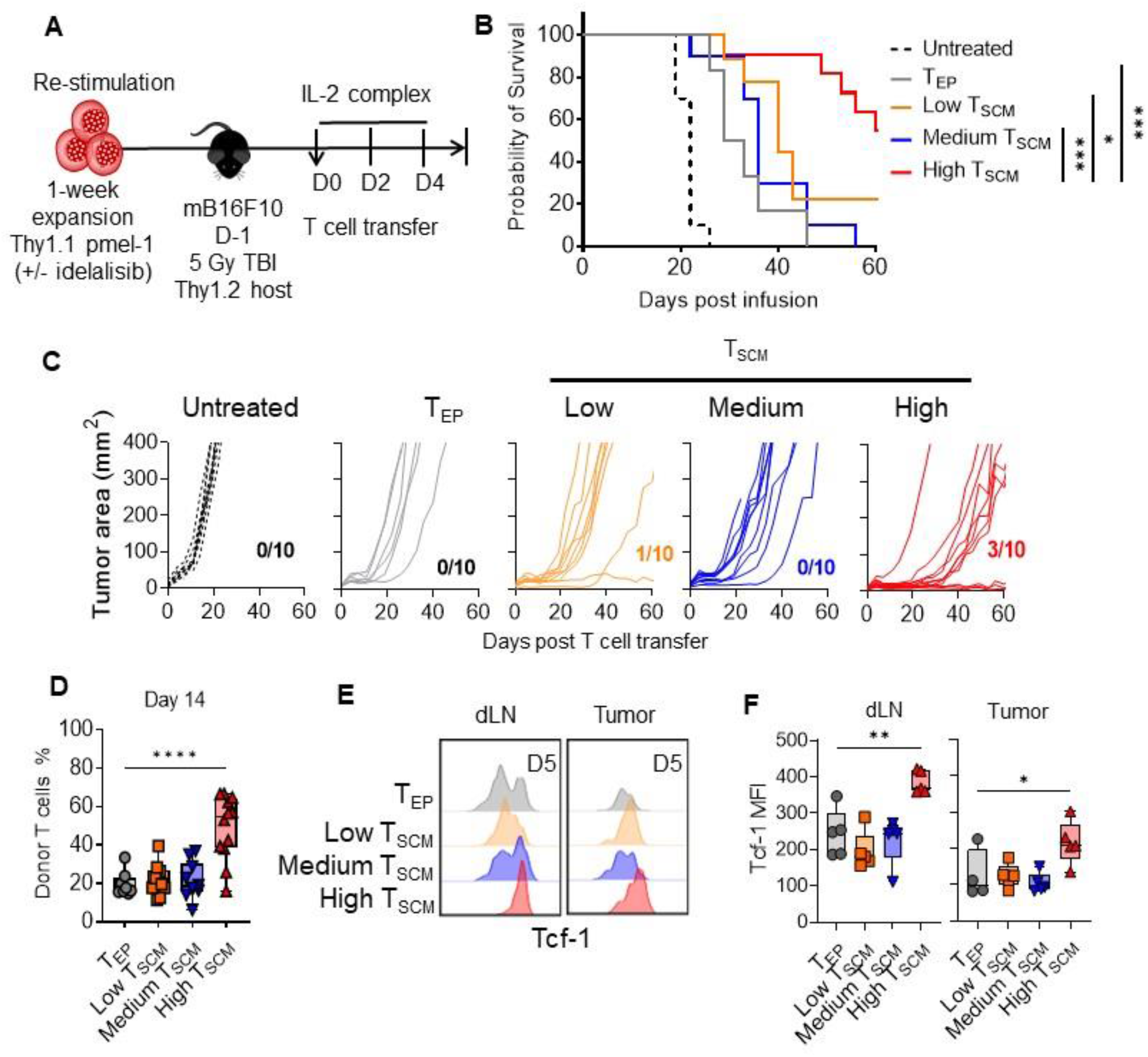
High T_SCM_ cells mediate prolonged antitumor responses and sustain Tcf-1 *in vivo* compared to lower T cell stemness states. **A**) Cell therapy schema: adoptive transfer of various pmel-1 T cells into mice bearing poorly immunogenic melanoma. C57BL6 mice were injected subcutaneously with 5×10^5^ B16F10 cells and allowed to grow to ~100mm^2^. Lymphodepletion at 5Gy total body irradiation was performed the day prior T cell infusion. Five million pmel-1 T cells of each subset were then transferred, and IL-2 complex was given intraperitoneally every 2 days for three doses. Survival and tumor growth my measuring area over time was tracked. **B**) Survival curves of mice using Mantel-cox analysis show statistically significant differences between T_EP_ and T_SCM_ states in ACT. **C**) Single curves of tumor growth areas and surviving mice denoted per group. N=10 mice per group, representative of 3 independent experiments. **D**) T cell engraftment in the peripheral blood at day 14 between T_SCM_ and T_EP_ groups. N=8-15 per group, representative of 3 independent experiments. **E**) Flow cytometry histogram for Tcf-1 expression between draining lymph node (dLN) and tumor at day 5 following ACT. **F**) Quantitation of MFI of Tcf-1 in T_EP_ and T_SCM_ subsets within dLN or tumor. N=3-6 per group, representative of 2 independent experiments. ANOVA analysis with Bonferroni post-hoc correction for multiple comparisons. * p≤0.05, ** p≤0.01, *** p≤0.001, **** p≤0.0001 were considered significant, while ρ values >0.05 were considered not significant (ns).

### Long-term protection against melanoma recurrence is mediated by high T_SCM_

Low dose (~0.1μM) idelalisib was reported to enrich T cells with stemness, and mediate immunity to hematological malignancies (*37*, *38*). However, we show herein that low and medium T_SCM_ did not improve immunity to melanoma compared to high T_SCM_ cells (**Fig. 4B-C**). Yet, low T_SCM_ cells existed in a less differentiated state and expressed Tcf1 *in vitro*. Thus, posited that low T_SCM_ might mediate improved primary antitumor responses and even durable protective immunity against a more immunogenic melanoma tumor. To test this idea, as in **Fig. 5A**, we performed ACT in mice bearing a B16F10 melanoma tumor containing a three amino acid residue change at the N terminus in the mouse gp100. The residues in gp100 were converted from EGS to KVP, for which the TCR of the pmel-1 T cells have a greater affinity and was reported to behave similar to a tumor “neoantigen” (*39*). Indeed, low, medium, and high T_SCM_ were equally effective at mediating primary immune responses to KVP melanoma (**Fig. 5B**). Yet most mice treated with T_EP_ cells died within one month. At day 50, survivor mice in all groups were given a second subcutaneous rechallenge with KVP B16F10. Mice infused with low and medium T_SCM_ eventually relapsed post tumor rechallenge. Mice protected after the second rechallenge were then given third subcutaneous tumor with the poorly immunogenic parental EGS B16F10 on day 80 (**Fig. 5A-B**). After this challenge, more mice treated with low T_SCM_ cells relapsed. Finally, to test if mice in **Fig. 5B** that survived multiple subcutaneous melanomas rechallenges were protected against disseminated disease to the lungs, B16F10 melanoma was administered intravenously, and then allowed to grow for 21 days (**Fig. 5C**). As expected, metastatic tumor lesions were abundant in mice that never received T cell transfer compared to mice treated with TEP, low or medium TSCM cells (**Fig. 5D**). Strikingly, lung metastasis were less prevalent in animals originally infused with high T_SCM_ cells (**Fig. 5E**). Remarkably, tissue distribution analysis showed that >120 days following initial infusion, high T_SCM_ were enriched in the draining LN and in the lung (**Fig. 5F-G**).

**Figure 5.**
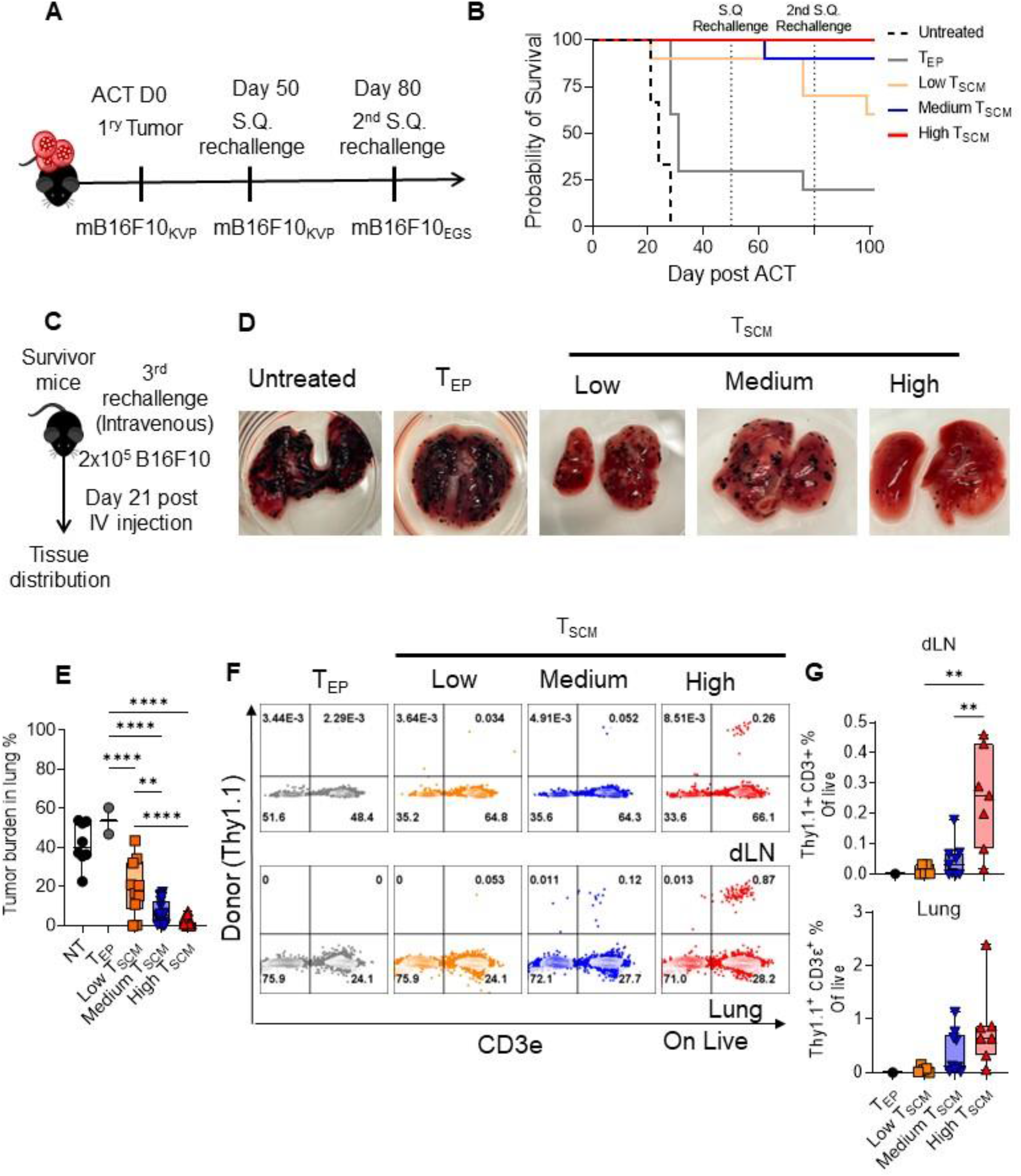
Long-term protection against immunogenetic melanoma is optimally mediated by high T_SCM_ cells. **A**) Multiple rechallenges to cell therapy schema: following primary tumor clearance by ACT with various T subsets (as in Fig 4A) in mice bearing “neoantigen” KVP B16F10 melanoma tumors, mice were given a second subcutaneous tumor challenge on the opposite flank at day 50. If mice continued to survive after this second rechallenge, a third parental B16F10 tumor injection was SQ implanted in the abdomen of mice on day 80. **B**) Survival of mice given this multiple rechallenges were tracked following ACT. N=10 mice per group representative of 2 independent experiments. **C**) Mice that survived a third subcutaneous rechallenge were then given 2.0×10^5^ B16F10 tumor cells through the tail vein on day 100 post ACT. Tumors were allowed to grow for 21 days and then animals were sacrificed and lung and tumor draining lymph node was harvested. **D**) Representative pictures of tumor laden lung of mice given T cells with different grades of stemness following multiple tumor challenge. N=2-10 mice per group. **E**) Quantitation of tumor burden by percentage of total lung area per lung lobe. N=2-10 mice per group. **F**) Flow plots of live transferred (Thy1.1) T cells in either lung or draining lymph node of original tumor site. **G**) Quantitation of T cell engraftment in draining lymph node from primary tumor and lung denoted by congenic mismatch, donor (Thy1.1^+^ CD3^+^) T cells. ANOVA analysis with Bonferroni post-hoc correction for multiple comparisons. * p≤0.05, ** p≤0.01, *** p≤0.001, **** p≤0.0001 were considered significant, while ρ values >0.05 were considered not significant (ns).

### PI3Kδ signaling blockade generates human TIL and CAR T cell with differential stemness levels

For translational relevance to human adoptive T cell therapies (TIL or CAR T cells), we next evaluated whether degrees of stemness induced by PI3Kδ inhibition could be achieved in T cells that infiltrate human tumors (TIL). To test this idea, metastatic tumors from draining lymph nodes of patients with melanoma or Merkel cell carcinoma were expanded as in **Fig. 6A**. Specifically, human melanoma and Merkel cell carcinoma metastatic lesions from draining lymph nodes were cultured with CD3 agonist and 6,000IU/mL of IL-2 for 5 days, once T cells grew from lesions, cultures were treated with escalating idelalisib doses (DMSO, 0.1μM, 1μM or 10μM) for 10-14 days. Consistent with our findings from murine T cells, both CD8^+^ and CD4^+^ from patient TILs exposed to high dose PI3Kδ inhibition expressed high levels of IL-7Ra but less granzyme B compared to TIL treated with lower dose PI3Kδ inhibitor (**Fig. 6B-C**). Further, Tcf-1 and Lef-1 were most enriched in TIL treated with high dose idelalisib compared to those given lower doses of this drug (**Fig. 6D-E**). To test if human T cells with differing stemness levels could mediate anti-tumor activity, we engineered them with a second-generation CAR construct containing a CD3ζ signaling domain, a 4-1BB co-stimulatory domain, and a mesothelin-specific scFv antibody region. Mesothelin specific CAR T cells were expanded with CD3/CD28 activation beads in the presence of low (0.1μM), medium (1μM) or high concentrations of idelalisib (10μM), expanded for a week in the presence of 200IU/mL IL-2 and then infused into NSG mice bearing M108 mesothelioma tumors. High CAR T_SCM_ cells best controlled tumor growth compared to low or medium CAR T_SCM_ cells (**Fig. 6F**). As anticipated, T_EP_ CAR T cells were largely ineffective at regressing human mesothelioma tumors. Collectively, these findings suggest that the stemness features observed in our murine models can be recapitulated *in vitro* in both human TIL and CAR T cells. Importantly T cells with the most “stemness” possess the potential to regress human tumors more effectively in relevant tumor models.

**Figure 6.**
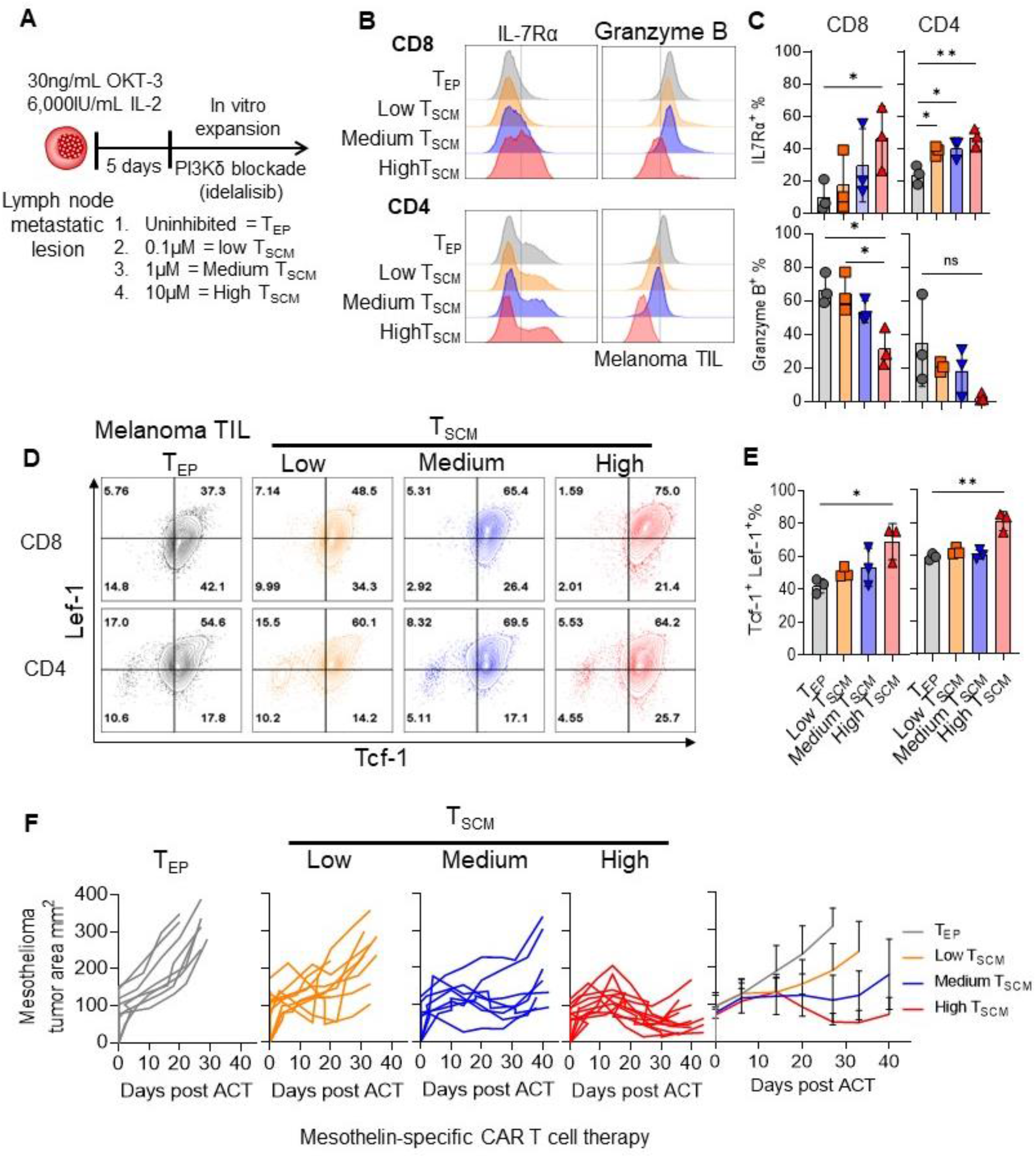
Progressive PI3K*δ* blockade enriches human CD4^+^ and CD8^+^ TILs and CAR T cells with graded stemness and distinct tumor responses. **A**) Human melanoma or Merkel cell carcinoma metastatic lesions to draining lymph nodes were cultured with CD3 agonist and 6,000IU/mL of IL-2 for 5 days, once T cells grew from lesions, cultures were treated with escalating doses of idelalisib (0.1μM, 1μM and 10μM) for 10-14 days prior to analysis. Nomenclature previously established for T_SCM_ grades. **B**) TIL expanded for ~14-20 days with idelalisib was assessed for IL7Rα and Granzyme B. **C**) Quantitation of **B** positive populations in CD4 and CD8 TIL. **D**) Tcf-1 and Lef-1 in TIL subsets. **E**) Quantitation of Tcf-1^+^ Lef-1^+^ populations in CD4 and CD8 TIL. N=3 patient samples, 2 melanoma, 1 Merkel cell carcinoma. **F**) single and summary tumor growth curves of human M108 mesothelioma over time after transfer of bulk CAR T cells (2×10^6^/mouse) transduced with a mesothelin specific CAR, expanded with low (0.1μM), medium (1μM) or high (10μM) concentrations of idelalisib and subsequently transferred into mesothelioma harboring NSG mice. N=8-10 per group. Representative of 2 independent experiments. ANOVA analysis with Bonferroni post-hoc correction for multiple comparisons. * p≤0.05, ** p≤0.01, *** p≤0.001, **** p≤0.0001 were considered significant, while ρ values >0.05 were considered not significant (ns).

### High T_SCM_ mediate durable anti-tumor immunity in a Tcf-1 and Lef-1 dependent manner

Tcf-1 has been a well-defined transcription factor in TIL that is correlated with enhanced survival in patients, however Lef-1 is also an important but underexplored transcription factor involved in stemness and is associated with Tcf-1 function (*40*, *41*). To test whether Tcf-1 or Lef-1 are equally important for the success of ACT, we used CRISPR to knock out Tcf-1 or Lef-1 in pmel-1 high T_SCM_ cells, resulting in knockout efficiencies of ~65% and ~75% in Tcf-1 and Lef-1 respectively (**Fig. 7A-B**). As expected, high T_SCM_ anti-tumor activity and survival was impaired when Tcf-1 was knocked out. Interestingly, however, Lef-1 was equally important in bolstering immunity, as its partial depletion also impaired the efficacy of adoptively transferred T_SCM_ cells (**Fig. 7C**), resulting in impaired survival in mice (**Fig. 7D**) even though they expressed Tcf-1 *in vitro* (**Fig. 7A-B**). Indeed, our work shows for the first time a nonredundant requirement for Lef-1 and Tcf-1 in the effectiveness of T_SCM_ mediated anti-tumor activity.

**Figure 7.**
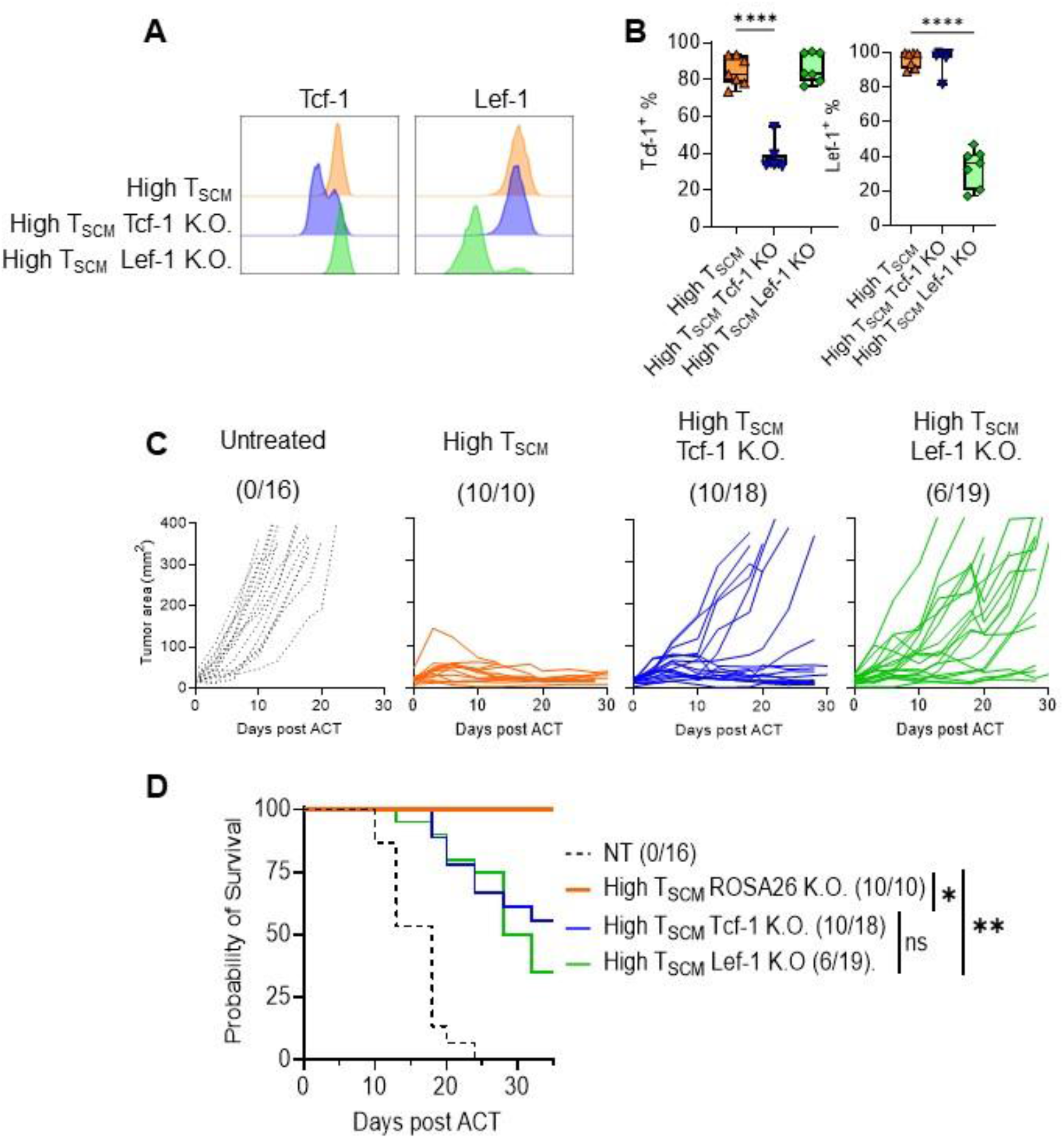
Tcf-1 and Lef-1 are both required for the sustained anti-tumor immunity of high T_SCM_ cells. **A**) CRIPSR knockout of Tcf-1 and Lef-1 representative histogram of pmel-1 high T_SCM_ cells cultured after 72hr following peptide stimulation. **B**) Quantitation of knocked out populations for Tcf-1 and Lef-1. N=7 per group, representative of 3 independent experiments. **C**) Tumor growth area over time measured by calipers for untreated, *high* T_SCM_ and high T_SCM_ knocked out of Tcf-1 or Lef-1. **D**) Survival proportions of mice from untreated, *high* TSCM and high TSCM knocked out of Tcf-1 or Lef-1. Mantel-Cox statistical test between groups. n=5-10 per condition, representative of 3 independent experiments. ANOVA analysis with Bonferroni post-hoc correction for multiple comparisons. ρ≤0.05, * p≤0.05, ** p≤0.01, *** p≤0.001, **** p≤0.0001 were considered significant, while ρ values >0.05 were considered not significant (ns).

## Discussion

Clinical responses have been observed in chemotherapy and immunotherapy refractory melanoma patients following adoptive T cell therapy (ACT) (*1*). However, limited efficacy to ACT can occur in patients and is frequently attributed to poor quality T cells, which are either terminally differentiated, have diminished host engraftment and/or compromised bioenergetics (*41*). The molecular cues regulating these processes are mediated by signals downstream of the PI3Kδ pathway, which are induced by the T cell receptor, costimulatory molecules, and cytokines (*42*–*44*). Our data suggest that T cell stemness grades can be generated *in vitro* through repurposing FDA-approved inhibitors of PI3Kδ. We further demonstrate that adequately characterizing metabolic and functional properties as a complement to phenotypic and transcriptional signatures is warranted to ascertain the adequate level of stemness for expansion of highly therapeutic ACT products. We find that tumor specific T cells with high stemness features express T-bet, Tcf-1 and Lef-1 while suppressing Eomes, granzyme and TIM-3 expression, even after re-encounter of tumor antigen. Because of the overlap between memory T cell subsets and our *in vitro* generated stem-like memory T cells, identifying features that discern between suboptimal and optimally generated TSCM for ACT remains a possible challenge. Additionally, exhaustive characterization of T cells for adoptive immunotherapy can delay and add to the cost of the generation of ACT products.

We demonstrate that select phenotypic and functional characteristics of highly effective tumor specific stem-like memory T cells can help differentiate between poor and highly effective T_SCM_. Transcriptionally, *Plagl1* is a particular gene of interest, as our transcriptional data shows it is selectively different in T cells with high features of stemness, compared to those with poor tumor control. *Plagl1* is a regulatory gene in the Wnt/Catenin signaling pathway and is also selectively enriched in T cells that respond to PD-1 blockade in models of chronic viral infection (*10*). Another marker that can help delineate potent T_SCM_ is Bcl-2, as our data shows that only highly effective T_SCM_ express it at high levels before tumor antigen stress. In fact, Bcl-2 overexpression in CAR T cells can modestly enhance anti-tumor regression in other models (*45*). Further, we identified the prevention of exhaustion marker acquisition (such as Tim-3 and CD39) and preservation of mitochondrial functionality as potential markers to identify therapeutic stem-like T cells for ACT. Our data indicates that T cells with high degrees of stemness can display distinct bioenergetic features. Namely, T cells with high features of stemness diminish glucose use during primary expansion, but upon antigen re-encounter they greatly induce glycolytic genes with sustained mitochondrial mass and function.

Finally, we found that T_SCM_ with high levels of stemness sustained Tcf-1 and Lef-1 in the tumor and draining lymph node *in vivo* 5 days after initial infusion while low and medium T_SCM_ did not. We reveal that the anti-tumor activity of these cells requires both Tcf-1 or Lef-1, as knocking out either factor blunted their potency. Although Tcf-1 is likely required for the anti-tumor response of T cells, whether Lef-1 is important for tumor immunity has been less described (*34*, *46*, *47*). Lef-1 is important for the development and memory recall of CD8^+^ T cells during acute and chronic viral infection. Although, in these studies Tcf-1 played a more critical role than Lef-1 in the biology of mature T cells. (*48*, *49*). In fact, it appears that Lef-1 plays non-overlapping functions in effector T cells compared to Tcf-1, as with conditional deletion in solid tumors (ovarian, melanoma and neuroblastoma), specific effector T cells show impaired primary tumor response (*50*). Here we demonstrate, in contrast to effector T cells, that in T cells with high levels of stemness, Tcf-1 and Lef-1 are required to mediate potent and long-lived tumor immunity. Lef-1 and Tcf-1 have overlapping and redundant functions during T cell development; however, recent studies indicate that Tcf-1 and Lef-1 may have non-redundant and equally important roles in anti-tumor T cell immunity. Whether T cell memory subtypes generated between viral infections or tumors differ in their reliance for Lef-1 is a remaining question raised by our work, as previous work had shown Tcf-1 as a primary requirement for T cell mediated immunity of chronic viral infections. In conclusion, our findings shed light on a practical and translatable way to effectively and easily enrich potent stem-like memory T cells *in vitro* for both TIL and CAR T cell expansion using currently FDA-approved small molecule inhibitors of PI3Kδ and uncover the importance of Tcf-1 and Lef-1 in stem-like memory T cells with protective immunity to large established solid tumors.

## Acknowledgements

Research reported in this publication was supported in part by the Cancer Tissue and Pathology shared resource of Winship Cancer Institute of Emory University and NIH/NI (P30CA138292), Pediatrics/Winship Flow Cytometry Core Children’s Healthcare of Atlanta and NIH/NCI (P30CA138292), and flow Cytometry and Cell Sorting Shared Resource at Hollings Cancer Center, Medical University of South Carolina (P30 CA138313). This work was also supported by Melanoma Research Foundation (to G.O Rangel Rivera, H.M. Knochelmann), ARCs Foundation (to A.C. Cole), NCI F30 CA243307, NIH T32 GM008716 and DE017551, and; NIH T32 AI132164-01 and T32 DE017551 (to C.J. Dwyer); NCI F31 CA232646-01A1 and Hollings Cancer Center Graduate Fellowship (to A.S. Smith); NIH R50 CA233186 (to M.M. Wyatt); NCI R01CA248359-01 (to J.E. Thaxton), NCI R01CA228406, R21CA266088, P30CA138292 (to G.B. Lesinski), NCI R01 CA175061, R01 CA208514 plus MUSC and Emory University Start Up Funds (to C.M. Paulos). We would like to thank Drs. Nicholas Restifo and Carl H. June for sharing of materials.

