## Supplemental figures for "The degree of T cell stemness differentially impacts the potency of adoptive cancer immunotherapy in a Lef-1 and Tcf-1 dependent manner"

### Supplementary figures and legends

One the following pages, this document contains three supplemental figures related to the main document. Supplemental figure 1 and 2 are relevant to main figure 1.

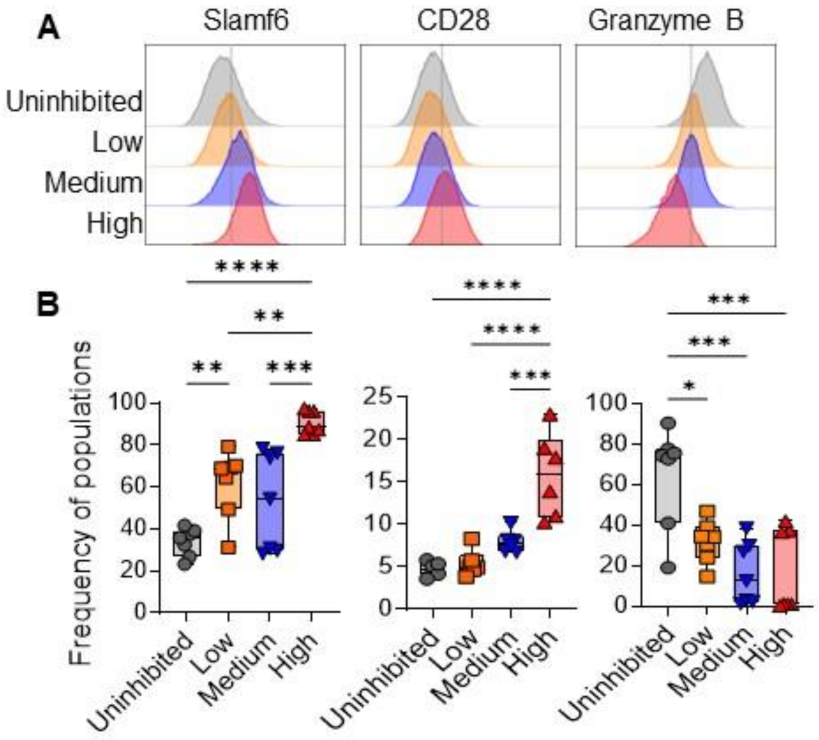


**Supplemental Figure 1**. **PI3Kδ inhibition enriches stem-like memory T cell phenotype. A**) Slamf6, CD28 and granzyme B after primary expansion with increasing idelalisib. **B**) Quantitation of positive populations for each marker on A. N=6 per condition, representative of 3 independent experiments. ANOVA analysis with Bonferroni post-hoc correction for multiple comparisons. $\rho \leq 0.05$, * $p \leq 0.05$, ** $p \leq 0.01$, *** $p \leq 0.001$, **** $p \leq 0.0001$ were considered significant, while $\rho$ values >0.05 were considered not significant (ns).

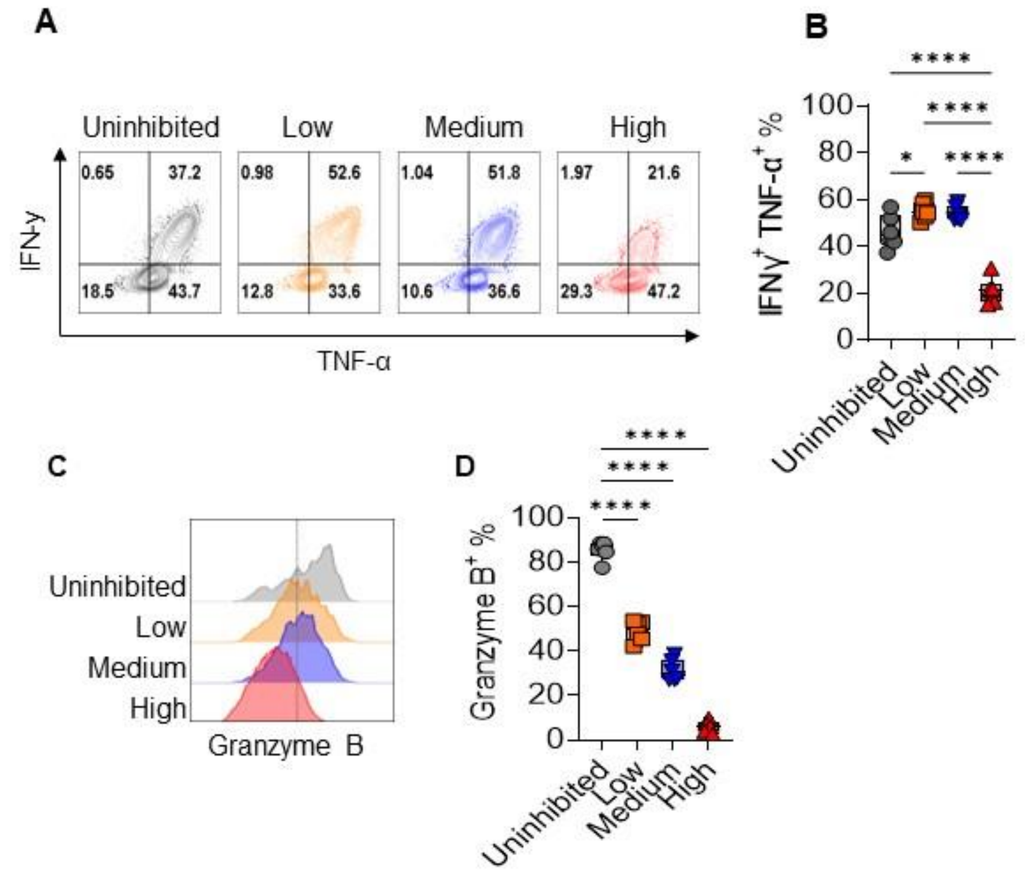


**Supplemental Figure 2**. **PI3Kδ inhibition blunts cytokine production and suppresses granzyme.** **A**) Interferon γ and TNF-α co-expression following antigen rechallenge for 24hr. **B**) Quantitation of double positive populations in H. N=6 per group, representative of 3 independent experiments. **C**) Granzyme B after 24hr antigen rechallenge, representative histogram. **D**) Quantitation of Granzyme positive populations. N=6 per group, representative of 4 independent experiments. ANOVA analysis with Bonferroni post-hoc correction for multiple comparisons. $\rho \leq 0.05$, * $p \leq 0.05$, ** $p \leq 0.01$, *** $p \leq 0.001$, **** $p \leq 0.0001$ were considered significant, while $\rho$ values $>0.05$ were considered not significant (ns).
