## Supplemental table 1 for "The degree of T cell stemness differentially impacts the potency of adoptive cancer immunotherapy in a Lef-1 and Tcf-1 dependent manner"

**Supplemental tables**

Reagent lists

**Mouse antibodies for flow cytometry**

| Antibody | Fluorophore | Clone | Vendor | Catalogue no |
|---|---|---|---|---|
| CD26 | FITC | H194-112 | Biolegend | 137805 |
| Eomes | PE-eFluor610 | WD1928 | Thermo Fischer | 61-4877-42 |
| T-bet | PerCP-Cy5.5 | 4B10 | Biolegend | 644805 |
| Lef-1 | AF647 | C12A5 | Cell signaling | 14022s |
| Granzyme B | Af700 | QA16A02 | Biolegend | 372221 |
| PD-1 | APC Cy7 | 29F.1A12 | Biolegend | 35223 |
| TIM3 | BV421 | RMT3-23 | Biolegend | 119723 |
| CD44 | BV570 | IM7 | Biolegend | 103037 |
| CD62L | BV605 | MEL-14 | Biolegend | 104437 |
| CD69 | BV650 | H1.2F3 | Biolegend | 104541 |
| LAG-3 | BV711 | C9B7W | Biolegend | 125243 |
| CD4 | BV750 | GK1.5 | Biolegend | 100467 |
| CD44 | BV786 | IM7 | Biolegend | 103041 |
| CD3e | BUV395 | 145-2C11 | BD | 565992 |
| Zombie UV | BUV515 | | Biolegend | 423107 |
| CD8a | BUV737 | 53-6.7 | BD | 612759 |
| SLAMF6 | BUV805 | 13G3 (RUO) | BD Optibuild | 748564 |
| CD25 | PE | 3C7 | Biolegend | 101903 |
| Tcf-1 | PE-Cy7 | C63D9 | Cell signaling | 90511 |
| Ly-6C | APC-Cy7 | HK1.4 | Biolegend | 128015 |
| CD28 | ACP | 37.51 | Biolegend | 102109 |
| SLAMF6 | BUV805 | 13G3 | BD Bioscience | 748564 |
| IL7Ra | BV421 | A7R34 | Biolegend | 135023 |
| CD62L | APC | MEL-14 | Biolegend | 104411 |
| CD62L | PE-Cy7 | MEL-14 | Biolegend | 104417 |
| CD62L | BV650 | MEL-14 | Biolegend | 104453 |
| CD44 | PrcPCy5.5 | IM7 | Biolegend | 103031 |
| T-bet | BV711 | 4B10 | Biolegend | 644819 |
| CD39 | PE | Duha59 | Biolegend | 143803 |
| IFN-γ | BV421 | XMG1.2 | Biolegend | 505829 |
| IFN-γ | APC-Cy7 | XMG1.2 | Biolegend | 505849 |
| TNF-α | PE-Cy7 | MP6-XT22 | Biolegend | 506323 |
| PI | | | Biolegend | 423301 |
| Annexin V | APC | | Biolegend | 640919 |

**Human antibodies for flow cytometry**

| Antibody | Fluorophore | Clone | Vendor | Catalogue no |
|---|---|---|---|---|
| Lef-1 | AF647 | C12A5 | Cell signaling | 14022s |
| Granzyme B | Af700 | QA16A02 | Biolegend | 372221 |
| Zombie UV | BUV515 | | Biolegend | 423107 |

| | | | | |
|---|---|---|---|---|
| Tcf-1 | PE-Cy7 | C63D9 | Cell signaling | 90511 |
| CD4 | BUV395 | RPA-T4 | BD Bioscience | 564724 |
| CD8 | BV605 | SK1 | Biolegend | 344741 |
| CD3ε | FITC | OKT3 | Biolegend | 317305 |
| IL-7RA | BV421 | A019D5 | Biolegend | 351309 |

**Western antibodies**

| Target | Host | Clone | Vendor | Catalogue no | Dilution |
|---|---|---|---|---|---|
| Actin-β | Rabbit | 13E5 | Cell Signaling | 4970S | 1:2000 |
| α-Tubulin | Mouse | DM1A | Cell Signaling | 3873T | 1:1000 |
| BCL-2 | Rabbit | D17C4 | Cell Signaling | 3498T | 1:1000 |
| Phospho-Akt (Ser473) | Rabbit | Ser473 | Cell Signaling | 9271T | 1:1000 |
| Phospho-Akt (Thr308) | Rabbit | 244F9 | Cell Signaling | 4056S | 1:1000 |
| Akt (pan) | Rabbit | 11E7 | Cell signaling | 4685S | 1:1000 |
| Phospho-GSK-3α/β (Ser21/9) | Rabbit | Ser21/9 | Cell Signaling | 9331S | 1:1000 |
| Total GSK-3α/β | Rabbit | 27C10 | Cell Signaling | 9315S | 1:100 |
| Phospho-S6 Ribosomal Protein (Ser235/236) | Rabbit | D57.2.2E | Cell Signaling | 2211S | 1:1000 |
| Total S6 Ribosomal Protein | Rabbit | 5G10 | Cell Signaling | 2217S | 1:1000 |
| Phospho-4E-BP1 (Thr37/46) | Rabbit | 236B4 | Cell Signaling | 2855S | 1:1000 |
| Total 4E-BP1 | Rabbit | 53H11 | Cell Signaling | 9644S | 1:1000 |
| Phospho-mTOR (Ser2448) | Rabbit | Ser2448 | Cell Signaling | 2971S | 1:1000 |
| Total mTOR | Rabbit | 7C10 | Cell Signaling | 2983S | 1:1000 |

**Dyes for flow cytometry**

| Reagent | Vendor | Catalogue number | Final concentration |
|---|---|---|---|
| Fixation buffer | Biolegend | 420801 | Per manufacturer |
| Foxp3 /Transcription Factor Fixation/Permeabilization | eBioscience | 00-5521-00 | Per manufacturer |
| Zombie aqua | Biolegend | 423101 | 1:1000 |
| Zombie UV | Biolegend | 423107 | 1:1000 |
| TMRM | Thermo Fischer | T668 | 250nM |
| Mitotracker FM Deep Red | Thermo Fischer | M46753 | 20nM |
| 2-NBDG | Cayman | 600470 | 1:200 |

**Mice**

| Mouse | Backgrounds | Supplier | RRID |
|---|---|---|---|
| NOD.Cg-Prkdcscid Il2rgtm1Wjl/SzJ | NOD/ShiLtJ | Jackson Laboratory | IMSR_JAX: 005557 |
| B6.Cg-Thy1a/Cy Tg(TcraTcrb)8Rest/J (Pmel-1) | C57/BL6N | Jackson Laboratory | IMSR_JAX:005023 |
| C57BL/6J (B6) | C57/BL6N | Jackson Laboratory | IMSR_JAX:000664 |

**Cell lines**

| Cell line | Backgrounds | Epitope | Supplier | RRID |
|---|---|---|---|---|
| B16F10 | C57/BL6N | Mouse gp100 (EGS) | ATCC | CVCL_0159 |
| B16F10 hgp100 | C57/BL6N | Human (KVP) and mouse gp100 (EGS) | Gift from Restifo Lab | N/A |
| M108 | Mesothelioma patient derived | Mesothelin | Gift from June Lab | N/A |

**Inhibitors**

| Reagent | Vendor | Catalogue number | Final concentration |
|---|---|---|---|
| Glucose | Sigma | D9434 | 10mM |
| 2-DG | Sigma | D6134 | 20mM |
| Oligomycin A | Sigma | 75351 | 1uM |
| FCCP | Sigma | C2920 | 1uM |
| Rotenone | Sigma | R8875 | 1uM |
| Antimycin A | Sigma | A8674 | 2uM |
| Idelalisib | Selleckchem | S2226 | As described |
| PMA | Sigma | P1585 | 300nM |
| Ionomycin | Sigma | I9657 | 200nM |
| Etomoxir Sodium salt | Cayman | 11969 | 3uM |

**qPCR reagents**

| Reagent | Vendor | Catalogue number |
|---|---|---|
| TRIzol™ Reagent | Thermo Fischer | 15596018 |
| Glycogen, molecular biology grade | Thermo Fischer | R0561 |
| iScript™ cDNA Synthesis Kit | Bio-Rad | 1708890 |
| SsoAdvanced Universal SYBR Green Supermix | Bio-Rad | 1725270 |

**Primer sequences**

| Target | 5'-3' sequence | Vendor |
|---|---|---|
| Tcf7 Forward | AGCATCCGCAGCCTCAAC | IDT |
| Tcf7 Reverse | GTGGACTGCTGAAATGTTCG | IDT |
| Lef1 Forward | CCCACACGGACAGTGACCTA | IDT |
| Lef1 Reverse | TGGGCTCCTGCTCCTTTCT | IDT |
| Actb Forward | ACGTAGCCATCCAGGCTGGTG | IDT |
| Actb Reverse | TGGCGTGAGGGAGAGCAT | IDT |
| 18s rna forward | CGC GGT TCT ATT TTG TTG GT | IDT |
| 18s rna reverse | AGT CGG CAT CGT TTA TGG TC | IDT |

**Seahorse reagents**

| Reagent | Vendor | Catalogue number |
|---|---|---|
| Seahorse XF Calibrant solution | Agilent | 103059-000 |
| Cell-Tak™ Cell and Tissue Adhesive | Corning | 35424 |
| Seahorse XF base medium, without phenol red | Agilent | 103335-100 |
| Seahorse XF 100 mM pyruvate solution | Agilent | 103578-100 |
| Seahorse XF 200 mM glutamine solution | Agilent | 103579-100 |
| Seahorse XFe24 FluxPak cartridge | Agilent | 102342-100 |

**CRISPR reagents**

| Reagent | Vendor | Catalogue number |
|---|---|---|
| Alt-R® CRISPR-Cas9 tracrRNA, ATTO™ 550, 20 nmol | IDT | 1075928 |
| TrueCut 2.0 Cas9 | Thermo Fischer | A36498 |
| Neon™ Transfection System 100 µL Kit | Thermo Fischer | MPK10096 |
| Positive Control, Mouse Rosa26, mod-sgRNA | Synthego | N/A |

**CRISPR crRNA**

| crRNA guide | Gene | Position | Strand | Target sequence | PAM |
|---|---|---|---|---|---|
| Mm.Cas9.LEF1.1.AA | LEF1 | 131115496 | + | GCGACCCGTACATGTCAAAT | GGG |
| Mm.Cas9.LEF1.1.AD | LEF1 | 131111531 | + | ATGATCCCCTTCAAGGACGA | AGG |
| Mm.Cas9.LEF1.1.AE | LEF1 | 131113878 | + | TCAGGAGCCCTACCACGACA | AGG |
| Mm.Cas9.TCF7.1.AA | TCF7 | 52258890 | + | GAAGTGCTGTCTATATCCGC | AGG |
| Mm.Cas9.TCF7.1.AB | TCF7 | 52261556 | + | CTGCTGAAATGTTCGTAGAG | TGG |
| Mm.Cas9.TCF7.1.AC | TCF7 | 52257030 | + | TAAAGCATGAACGCATTGAG | GGG |
